# SARS-CoV-2 variant B.1.1.7 is susceptible to neutralizing antibodies elicited by ancestral Spike vaccines

**DOI:** 10.1101/2021.01.27.428516

**Authors:** Xiaoying Shen, Haili Tang, Charlene McDanal, Kshitij Wagh, Will Fischer, James Theiler, Hyejin Yoon, Dapeng Li, Barton F. Haynes, Kevin O. Sanders, Sandrasegaram Gnanakaran, Nick Hengartner, Rolando Pajon, Gale Smith, Filip Dubovsky, Gregory M. Glenn, Bette Korber, David C. Montefiori

**Affiliations:** Department of Surgery, Duke University School of Medicine, Durham, NC, USA; Duke Human Vaccine Institute, Duke University School of Medicine, Durham, NC, USA; Theoretical Biology and Biophysics, Los Alamos National Laboratory, Los Alamos, NM USA; Moderna Inc., Cambridge, MA, USA; Novavax, Inc., Gaithersburg, MD, USA; Department of Medicine, Duke University Medical Center, Durham, NC, USA

## Abstract

The SARS-CoV-2 Spike glycoprotein mediates virus entry and is a major target for neutralizing antibodies. All current vaccines are based on the ancestral Spike with the goal of generating a protective neutralizing antibody response. Several novel SARS-CoV-2 variants with multiple Spike mutations have emerged, and their rapid spread and potential for immune escape have raised concerns. One of these variants, first identified in the United Kingdom, B.1.1.7 (also called VUI202012/01), contains eight Spike mutations with potential to impact antibody therapy, vaccine efficacy and risk of reinfection. Here we employed a lentivirus-based pseudovirus assay to show that variant B.1.1.7 remains sensitive to neutralization, albeit at moderately reduced levels (~2-fold), by serum samples from convalescent individuals and recipients of two different vaccines based on ancestral Spike: mRNA-1273 (Moderna), and protein nanoparticle NVX-CoV2373 (Novavax). Some monoclonal antibodies to the receptor binding domain (RBD) of Spike were less effective against the variant while others were largely unaffected. These findings indicate that B.1.1.7 is not a neutralization escape variant that would be a major concern for current vaccines, or for an increased risk of reinfection.

## Introduction

Genetic evolution in the SARS-CoV-2 virus is an increasing concern for the COVID-19 pandemic. Continued high infection rates are providing opportunities for the virus to acquire mutations that contribute to virus spread and possible immune evasion. Mutations in the viral Spike are a particular concern because this glycoprotein mediates virus attachment and entry (1) and is the major target for neutralizing antibodies (2). The D614G spike variant that spread rapidly during March-April of 2020 (3, 4) and by June 2020 was found in most sequences globally, is the earliest evidence for adaptive evolution of this virus in humans. The D614G mutation imparts increased infectivity *in vitro* (5, 6), accelerated transmission in hamsters (6), and a modest increase in neutralization susceptibility (7), all of which are explained by a more open conformation of the receptor binding domain (RBD) (7, 8). The mutation does not appear to increase disease severity despite an association with higher virus loads in respiratory secretions (5). Notably, several vaccines proved highly efficacious in phase 3 trials conducted while D614G was the dominant variant in the global pandemic (9–11).

Newer variants with additional mutations are spreading rapidly in the United Kingdom (UK)(B.1.1.7), South Africa (501Y.V2) and Brazil (484K.V2) (12–14) (**Fig. S1**; for daily updates of the global sampling of these variants, see GISAID’s “Tracking of Variants” page https://www.gisaid.org/hcov19-variants/). Among them, the B.1.1.7 lineage of SARS-CoV-2 has caused public health concern because of it high rate of transmission in the UK (12). This variant, also called Variant of Concern 202012/01 (VOC 202012/01) (15), contains 17 non-synonymous mutations, including the D614G mutation and 8 additional mutations in Spike: ΔH69-V70, ΔY144, N501Y, A570D, P681H, T716I, S982A, and D1118H. Three B.1.1.7 Spike mutations were of particular concern: a two amino acid deletion at position 69-70 of the N-terminal domain (NTD); N501Y, located in the receptor binding motif (RBM); and P681H, proximal to the furin cleavage site (12). Each of these three mutations are also found in other variants of interest. Epidemiological evidence and mathematical modeling data suggest the variant is more transmissible than the SARS-CoV-2 variants that were circulating prior to its introduction (**Figure 1**) (16–19) and, though initially reported as not more pathogenic (20), evidence of increased mortality rate has also been reported (21). As mutations in Spike have potential to alter virus infectivity and/or susceptibility to neutralizing antibodies, one critical question is whether this B.1.1.7 variant will evade current vaccines, all of which are based on ancestral Spike.

**Figure 1.**
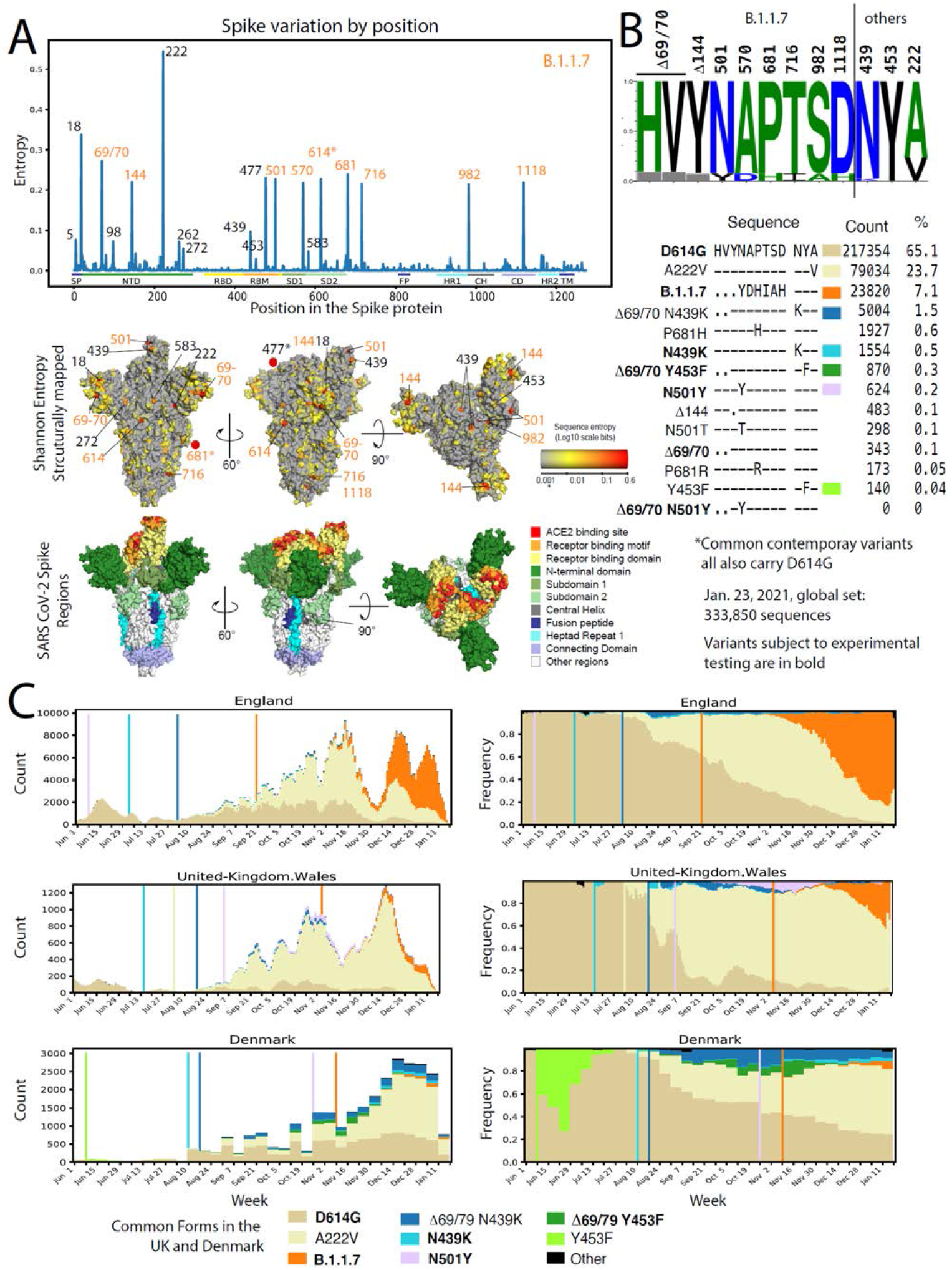
Epidemiology tracing of mutations in B.1.1.7 and co-circulating relevant mutations in the UK and Danish SARS-CoV-2 epidemics. **A**. Entropy scores summarizing the level of diversity found in positions in Spike. These scores are dependent on sampling, and recent sampling from the United Kingdom and Denmark has been particularly intense relative to other regions of the world (Fig. S1). B.1.1.7 mutations are highlighted in orange. The subset of B.1.1.7 sites with greater entropy scores (69/70, 681, and 501) are also often found in the context of other variants. The most variable site in Spike is at 222, and is indicative of the GV clade. G614 has dominated global sampling since June 2020, and the entropy at 614 reflects presence of the ancestral form, D614, sampled in the early months of the pandemic. These same entropy scores are first mapped by linear position in the protein and then mapped onto the Spike structure below the graph. Regions of Spike are indicated by the same colors in the linear and structural maps. **B**. Frequencies of variants in relevant positions. Using the Analyze Align tool at cov.lanl.gov, we extracted the columns of interest for the B.117 Spike mutations, and the additional sites of interest at 439, 453, and 222, out of a 333,850 sequence set extracted from GISAID on Jan 23, 2021. The LOGO at the top indicates the AA frequency in the full dataset; the grey boxes indicate deletions. All common forms of combinations of mutations at these sites of interest are shown, followed by their count and percentage. The forms that were common in the UK and Denmark are each assigned a color, and used to map transition in frequencies of these forms over time in part C. **C**. Weekly running averages for each of the major variants in the UK and Denmark, based on the variants shown in part B, are plotted; the actual counts are on the left, and relative frequencies on the right. Some windows in time are very poorly sampled, some very richly. The vertical lines indicate when a variant is first sampled in a region. Note the lavender N501Y in Wales; this is N501Y found out of the context of B.1.1.7 and transient. The shifts in relative prevalence from the G clade (beige, D614G), to the GV clade (cream, A222V), to the B.1.1.7 variants (orange).

Here we assessed the neutralization phenotype of the B.1.1.7 variant using convalescent sera, monoclonal antibodies (mAbs) and serum samples from phase 1 trials of an mRNA-based vaccine (mRNA-1273, Moderna) and a protein nanoparticle vaccine (NVX-CoV2373, Novavax). In addition, we characterized another two RBD mutations, N439K and F453Y that showed limited circulation in both Denmark and the UK preceding the circulation of the B.1.1.7 variant; these RBD mutations are each most often found coupled with the same ΔH69-V70 that is in B.1.1.7.

## RESULTS

### Rational for testing B.1.1.7 variant and select subvariants

B.1.1.7 contains 8 mutations in Spike (**Figure 1**), and the lineage is associated with many additional mutations throughout the SARS-CoV-2 genome (**Figure S2**). Among the Spike mutations, N501Y is suggested to increase ACE2-RBD interaction (22) and has been shown to be critical for adaptation of SARS-CoV-2 to infect mice (23). N501Y has twice reached frequencies between 10-20% in local populations as a single mutation in a D614G Spike backbone (once in Wales, (**Figure 1C** and **Figure S1**), and also once in Victoria, Australia) but in these cases it did not persist. N501Y is also evident in a distinctive variant that is increasing in frequency in South Africa, 501Y.V2, and accompanies other mutations in Spike that can confer partial resistance to convalescent sera (24, 25) and vaccine sera (25, 26). A double deletion of amino acids H69-V70 in the N-terminal domain (NTD) of Spike often co-occurs with one of 3 mutations in RBD: N501Y, N439K, or Y453F (27). Y453F is associated with a mink farm outbreak in Denmark, with and without the presence of a ΔH69/V70 deletion (27, 28), but is also found in people in Denmark and the UK (**Figure 1**). N439K mutation usually occurs with ΔH69/V70 but occurs frequently without the ΔH69/V70 mutation as well. In an *in vitro* selection study with Regeneron antibodies, Y453F and N439K were found to escape neutralization by REGN10933 (29) and REGN10987 (30) that comprise the REGN-COV2 cocktail regimen (31). N439K has also been reported to resist neutralization while maintaining virus fitness/infectivity (30, 32). Another mutation of obvious concern in B.1.1.7 is P681H, proximal to the furin cleavage site (**Fig. 1C, Fig. S2, and Fig. S3**) that has arisen many times independently (**Figure S1**) and has come to dominate the local epidemic in Hawaii.

### Neutralization of variant B.1.1.7 by serum from convalescent individuals and vaccine recipients

SARS-CoV-2 variant B.1.1.7 was compared to the D614G variant in neutralization assays with serum samples from 15 COVID-19 convalescent individuals, 40 recipients of the Moderna mRNA-1273 vaccine (11 samples from 29 days post first inoculation, Day 29; 29 samples from 28 days post second inoculation, Day 57) and 28 recipients of the Novavax Spike protein nanoparticle vaccine NVX-CoV2373 (2 weeks post second inoculation). Selection of NVX-CoV2373 vaccine serum samples was random and not pre-selected based on any selection criterion of anti-Spike or neutralizing titers. The B.1.1.7 variant was neutralized by all vaccine sera, although with modestly diminished susceptibility compared to the D614G variant (**Fig 2A, 2B**). A modest decrease in neutralization susceptibility also was seen with convalescent sera, although not to the same extent seen with vaccine sera. Median ID50 titers of sera from both phase 1 vaccine trials were on average 2.1-fold lower against B.1.1.7 than against D614G (**Table S1**). The fold difference in ID50 titer ranged from 0.36 to 8.62 for Moderna sera, with an interquartile range (IQR) of 1.6 to 2.9. The fold difference in ID50 titers ranged from 0.85 to >20 for Novavax sera, with an IQR of 1.5 to 3.0. Median ID80 titers of sera from both phase 1 trials were on average 1.7-fold lower against B.1.1.7 than against D614G (**Table S1**), with a tighter range of fold difference compared to ID50. The fold difference in ID80 ranged from 0.91 to 3.21 for Moderna sera, with an IQR of 1.4 to 1.9. The fold difference in ID80 titer ranged from 0.89 to 3.98 for Novavax, with an IQR of 1.5 to 2.6. Convalescent sera showed an average of 1.5-fold (group median) lower ID50 titer against the B.1.1.7 variant (range 0.7 to 5.5; IQR = 1.1 to 1.8) and 1.5-fold (group median) lower ID80 titer (range 0.7 to 3.3; IQR= 1.3 to 1.8). The fold differences were statistically significant with p <0.0001 for both ID50 and ID80 for Moderna and Novavax phase 1 sera, and p<0.001 for ID80 of both sets of vaccine sera and the convalescent sera (Wilcoxon signed-rank test, paired, 2-tailed; false discovery rate (FDR) corrected q values <0.1, corresponding to p <0.042 in this study, were considered as significant) (**Fig 2A, 2B, Table S2**).

**Figure 2.**
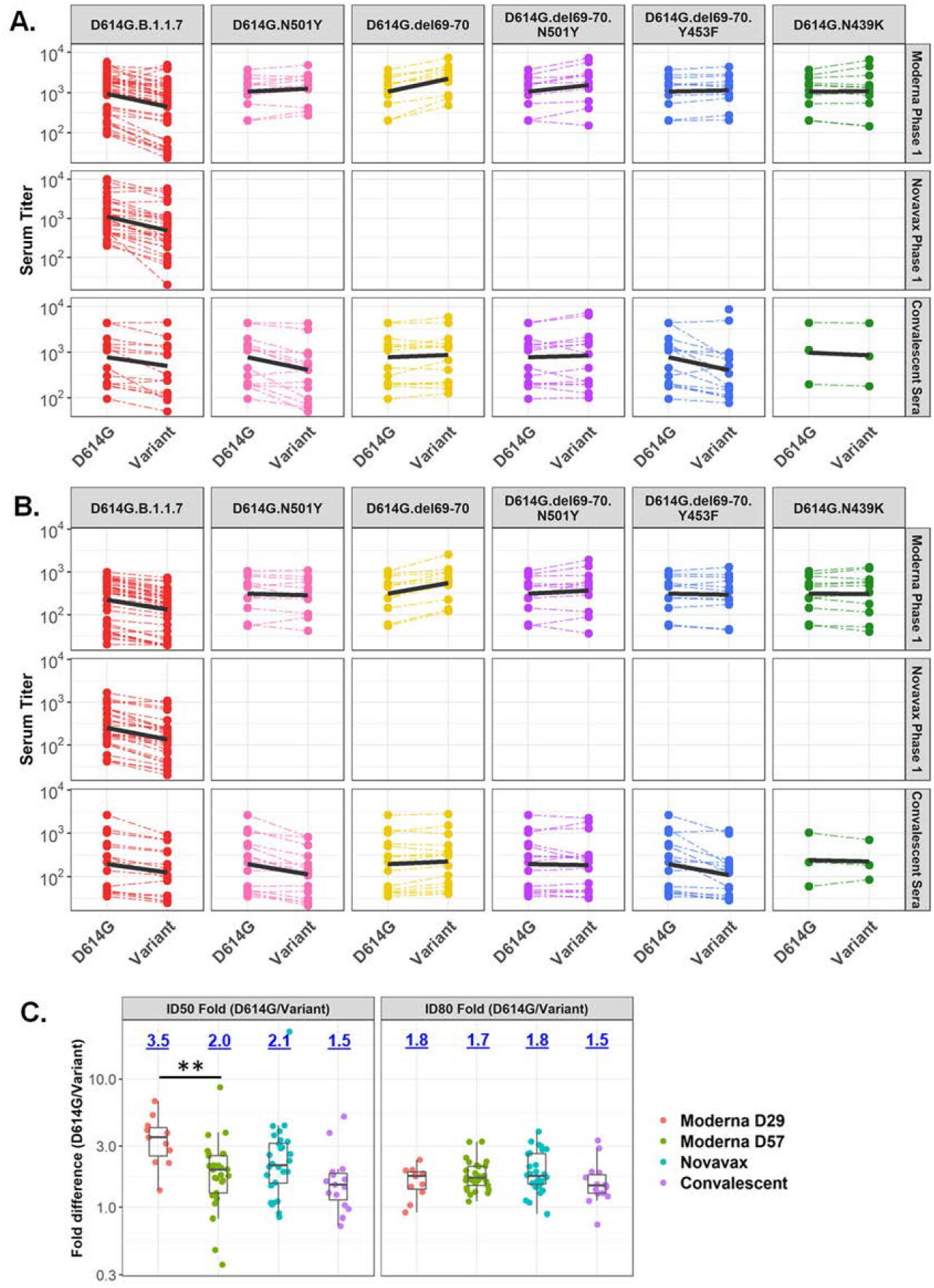
Neutralization of variants by vaccine and convalescent sera. Serum ID50 (**A**) and ID80 (**B**) titers of neutralization of each variant, as indicated on top of each panel, relative to D614G by vaccine sera (top 2 rows) and convalescent sera. Dashed thin lines represent individual samples, thick black lines represent geometric means of each sample group as indicated on the right. NT-not tested. Also see Table S1. (**C**) Fold decline of ID50 (left) and ID80 (right) titers for each variant over D614G (D614G/variant) for each serum sample set as identified. Numbers on top of each plot show median fold differences. Upper and lower border of each box represent IQR of the fold differences respectively, and the middle bars in boxes represent group median. Statistical significance of comparisons are indicated in all panels as: * p<0.042 (corresponding to q<0.1). ** p<0.01. *** p<0.001. **** p<0.0001. Wilcoxon signed-rank paired test for (A) and (B); Wilcoxon rank-sum test for (C). Also see Table S2.

Notably, sera with weaker neutralizing activity from the Moderna trial exhibited a more substantial reduction in activity against the B.1.1.7 variant. Because most low-titer samples in this trial were from Day 29 (single inoculation), we compared the change in neutralization titers for Day 29 and Day 57 samples from the Moderna trial as well as all samples from the Novavax trial and from convalescent individuals (**Fig 2C**). The Day 29 samples from the Moderna trial exhibited the greatest decrease in ID50 titer among the sample sets, and the decrease was significantly larger than the decrease for Day 57 samples against the B.1.1.7 variant (**Table S1**; p=0.0007), suggesting that antibody maturation can alleviate neutralization resistance.

### Neutralization of additional variants by serum from convalescent individuals and vaccine recipients

To gain insights into whether the reduced neutralization susceptibility of the B.1.1.7 variant was due to a single Spike mutation or a combination of two or more Spike mutations, we characterized B.1.1.7 subvariants containing either N501Y alone, ΔH69-V70 alone, and a combination of N501Y+ΔH69-V70, each in the D614G background. We also tested ΔH69-V70+Y453F and N439K in the D614G background because ΔH69-V70 is commonly shared with these mutations; they were first identified in association with recent outbreaks in minks and zoonotic transmission to humans in Denmark. Due to limited supplies, sera from the Novavax phase 1 trial were not included in these assays. Interestingly, the ΔH69-V70 mutation rendered the virus more susceptible to neutralization by Moderna (mRNA-1273 vaccine) sera but not convalescent sera (**Fig 2**). Median ID50 and ID80 titers for Moderna sera were 2-fold higher against D614G.ΔH69-V70 than against D614G (p<0.001) while titers of convalescent sera were comparable to D614G (**Fig 2A & 2B, Table S2**). N501Y had no significant impact on susceptibility to Moderna sera but did impart modest resistance to convalescent sera (**Fig 2**). Median ID50 and ID80 titers for convalescent sera against the N501Y variant were 2.2 and 1.8-fold lower (p<0.01 and p<0.001 for ID50 and ID80) while titers for Moderna sera were comparable to D614G (**Fig 2A & 2B; Table S2**). When both the ΔH69-V70 deletion and the N501Y mutation were present, the increase in susceptibility caused by ΔH69-V70 was diminished but still significant. No significant difference in neutralization titers was observed when both the ΔH69-V70 deletion and the N501Y mutation were present, except for a statistically significant (p<0.01) but minimal 1.18-fold increase in ID50 titer of Moderna sera against D614G.ΔH69-V70.N501Y compared to D614G (**Fig 2A & 2B, Table S2**). The variant with both the ΔH69-V70 deletion and the Y453F mutation showed decreased neutralization susceptibility to convalescent sera but not Moderna sera. Median ID50 and ID80 titers for convalescent sera were 1.7 and 1.5-fold lower against D614G.ΔH69-V70.Y453F than against D614G (p=0.012 [q=0.026] and p<0.001, respectively) (**Fig 2A & 2B, Table S2**). When neutralization titers for variants with and without the Y453F mutation were compared, median ID50 titers of Moderna and convalescent sera were 2.1-fold and 3.6-fold lower respectively, against D614G.ΔH69-V70.Y453F than against D614G.ΔH69-V70 (p<0.001 and p<0.0001 respectively) (**Table S2**). Median ID80 titers also were significantly lower for Moderna and convalescent sera against D614G.ΔH69-V70.Y453F than against D614G.ΔH69-V70 (1.8 and 2.5-fold; p<0.01 and p<0.001, respectively) (**Table S2**). Therefore, Y453F mutation can reverse the increased susceptibility conferred by the ΔH69-V70 mutation, demonstrating cooperative interactions between the RBD and NTD of Spike. The D614G.N439K variant showed neutralization titers comparable to D614G for Moderna sera (**Fig 2A & 2B, Table S2**).

### Neutralization by mAbs

Neutralization of the B.1.1.7 variant and corresponding subvariants was assessed with a panel of RBD-targeting mAbs (DH1041, DH1042, DH1043, DH1047, B38, H4, P2B-2F6, COVA1-18, COVA2-15, and S309; among with DH1047, COV1-18, and S309 are cross-reactive CoV RBD antibodies). The B.1.1.7 variant showed greatest resistance to mAbs B38, COVA2-15 and S309 (>10-fold difference in either IC50 or IC80 concentration compared to D614G) (**Table 1**). Resistance to B38 could not be fully explained by N501Y, ΔH69-V70 or the combination of these two mutations, whereas resistance to COVA2-15 and COVA1-18 was largely due to N501Y. Resistance to S309 was associated with N501Y, although this mutation alone accounted for only part of the resistance seen with the complete B.1.1.7 variant. The complete set of mutations and subsets of mutations in B.1.1.7 tested here had little (4.7-fold for DH1042 and H4 IC50) or no impact on other RBD antibodies (**Table 1**).

**Table 1.** Neutralization of variants by mAbs^a^.

| Virus | Value<sup>b</sup> | mAbs |  |  |  |  |  |  |  |  |  |
| --- | --- | --- | --- | --- | --- | --- | --- | --- | --- | --- | --- |
|  |  | B38 | COVA1-18 | COVA2-15 | DH1041 | DH1042 | DH1043 | DH1047 | H4 | P2B-2F6 | S309 |
| D614G | IC50 | 2.10 | 0.0060 | 0.0058 | 0.0094 | 0.011 | 0.0081 | 0.14 | 2.10 | 0.071 | 0.048 |
|  | IC80 | 12.00 | 0.031 | 0.028 | 0.036 | 0.068 | 0.040 | 0.54 | 18.00 | 1.20 | 0.69 |
| D614G.B117 | IC50 | **30**<br><u>(14.2x)</u> | 0.011<br>(1.8x) | **0.05**<br><u>(8.7x)</u> | 0.015<br>(1.6x) | **0.052**<br><u>(4.7x)</u> | 0.01<br>(1.3x) | 0.17<br>(1.2x) | **9.9**<br><u>(4.7x)</u> | 0.075<br>(1.1x) | **9.2**<br><u>(191.4x)</u> |
|  | IC80 | **>50**<br><u>(>4x)</u> | **0.16**<br><u>(5.1x)</u> | **0.95**<br><u>(34.5x)</u> | 0.039<br>(1.1x) | 0.13<br>(1.9x) | 0.05<br>(1.3x) | 0.64<br>(1.2x) | **>50**<br><u>(>3x)</u> | 2.2<br>(1.8x) | **>50**<br><u>(>72x)</u> |
| D614G.N501Y | IC50 | 4.7<br>(2.2x) | **0.032**<br><u>(5.2x)</u> | **0.084**<br><u>(14.6x)</u> | 0.01<br>(1.1x) | 0.012<br>(1.1x) | 0.015<br>(1.8x) | 0.072<br>(0.5x) | 3.7<br>(1.7x) | 0.043<br>(0.6x) | **0.15**<br><u>(3.1x)</u> |
|  | IC80 | **>50**<br><u>(>4x)</u> | **0.19**<br><u>(6.1x)</u> | **0.93**<br><u>(33.8x)</u> | 0.034<br>(1x) | 0.06<br>(0.9x) | 0.058<br>(1.5x) | 0.43<br>(0.8x) | 46<br>(2.5x) | 0.93<br>(0.8x) | **>50**<br><u>(>72x)</u> |
| D614G.del69-70 | IC50 | 1.5<br>(0.7x) | 0.0042<br>(0.7x) | 0.0051<br>(0.9x) | 0.007<br>(0.8x) | 0.0095<br>(0.9x) | 0.0077<br>(0.9x) | 0.1<br>(0.7x) | 2.5<br>(1.2x) | 0.1<br>(1.4x) | 0.018<br>(0.4x) |
|  | IC80 | 9.9<br>(0.8x) | 0.026<br>(0.8x) | 0.026<br>(0.9x) | 0.029<br>(0.8x) | 0.053<br>(0.8x) | 0.033<br>(0.8x) | 0.5<br>(0.9x) | 26<br>(1.4x) | 1.1<br>(0.8x) | **18**<br><u>(26.8x)</u> |
| D614G.del69-70.N501Y | IC50 | 2.5<br>(1.2x) | **0.026**<br><u>(4.3x)</u> | **0.051**<br><u>(8.8x)</u> | 0.0044<br>(0.5x) | 0.0061<br>(0.6x) | 0.0074<br>(0.9x) | 0.049<br>(0.4x) | 3.1<br>(1.5x) | 0.033<br>(0.5x) | 0.11<br>(2.2x) |
|  | IC80 | **40**<br><u>(3.3x)</u> | **0.18**<br><u>(5.8x)</u> | **1.6**<br><u>(56.4x)</u> | 0.025<br>(0.7x) | 0.039<br>(0.6x) | 0.035<br>(0.9x) | 0.28<br>(0.5x) | 30<br>(1.6x) | 1.0<br>(0.8x) | **>50**<br><u>(>72x)</u> |
| D614G.del69-70.Y453F | IC50 | 1.2<br>(0.5x) | 0.0083<br>(1.4x) | 0.0055<br>(0.9x) | 0.0046<br>(0.5x) | 0.004<br>(0.4x) | 0.011<br>(1.4x) | 0.048<br>(0.3x) | <u>0.64</u><br><u>(0.3x)</u> | 0.094<br>(1.3x) | 0.098<br>(2x) |
|  | IC80 | 15<br>(1.3x) | 0.043<br>(1.4x) | 0.03<br>(1.1x) | 0.027<br>(0.8x) | 0.03<br>(0.5x) | 0.05<br>(1.3x) | 0.24<br>(0.4x) | <u>5.7</u><br><u>(0.3x)</u> | 1.3 (1x) | **11**<br><u>(15.5x)</u> |
| D614G.N439K | IC50 | 1.3<br>(0.6x) | 0.0061<br>(1x) | 0.011<br>(1.9x) | 0.0075<br>(0.8x) | 0.0063<br>(0.6x) | 0.017<br>(2.1x) | <u>0.016</u><br><u>(0.1x)</u> | 3.6<br>(1.7x) | 0.15<br>(2.2x) | 0.046<br>(0.9x) |
|  | IC80 | 19<br>(1.6x) | 0.049<br>(1.6x) | **0.1**<br><u>(3.8x)</u> | 0.035<br>(1x) | 0.07<br>(1x) | 0.059<br>(1.5x) | 0.33<br>(0.6x) | 42<br>(2.3x) | 1.6<br>(1.3x) | 0.38<br>(0.6x) |

<sup>a</sup>Numbers in parenthesis followed by "x" represent fold differences over D614G.
<sup>b</sup>Unit of ID50 and ID80 concentrations is $\mu\text{g/mL}$ .
NA signifies data not available. Bold values: $>3$ -fold increase in ID50 or ID80 concentration over D614G, representing moderate decrease in susceptibility. Underlined and bold values: $>10$ -fold increase in ID50 or ID80 concentration over D614G, representing substantial decrease in susceptibility. Underlined: $<1/3$ the ID50 or ID80 of D614G, representing moderate increase in susceptibility.

The mAbs were largely unaffected by the Y453F and N439 mutations (**Table 1**). A modest increased sensitivity was seen in two cases: DH1047 assayed against D614G.N439K and H4 assayed against D614G.ΔH69-V70.Y453F. The latter variant also exhibited partial resistance to S309, which was mostly seen at IC80 and the level of change was similar to that caused by the ΔH69-V70 deletion alone, indicating the deletion is the cause of the decreased susceptibility rather than the Y453F.

We used structural analyses to understand the molecular mechanisms of mAb neutralization resistance or lack thereof. Mapping of the mAb epitopes (<4Å) on the Spike trimer showed that the RBD mutations were within, or close to the epitopes of all RBD antibodies, while ΔH69-V70 is in the NTD (**Figure 3**). N501Y and Y453F are in close proximity of the B38 paratope. Modeling of the N501Y mutation showed a potential clash between Y501 in Spike and S-30 in B38 light chain, consistent with the neutralization resistance of N501Y to B38. mAbs P2B-2F6, DH1041 and DH1043 had very similar epitopes and were grouped together (P2B-class in **Figure 3A**; also similar to Class 2 RBD mAbs (33). The RBD mutations were further from the P2B-2F6 and DH1041 paratopes, explaining the lack of impact of these mutations. Y453 is close (3.9Å) to the DH1043 paratope but with no predicted polar interactions. Structural modeling of N439K suggests a potentially more favorable interaction as Lys interacts with a negatively charged patch on the DH1047 surface. Although N501 is close to the DH1047 paratope, its side chain is oriented away from the mAb, suggesting no substantial impact due to N501Y. Y453 is also in close proximity of the B38 paratope but structural modeling showed no substantial impact.

**Figure 3.**
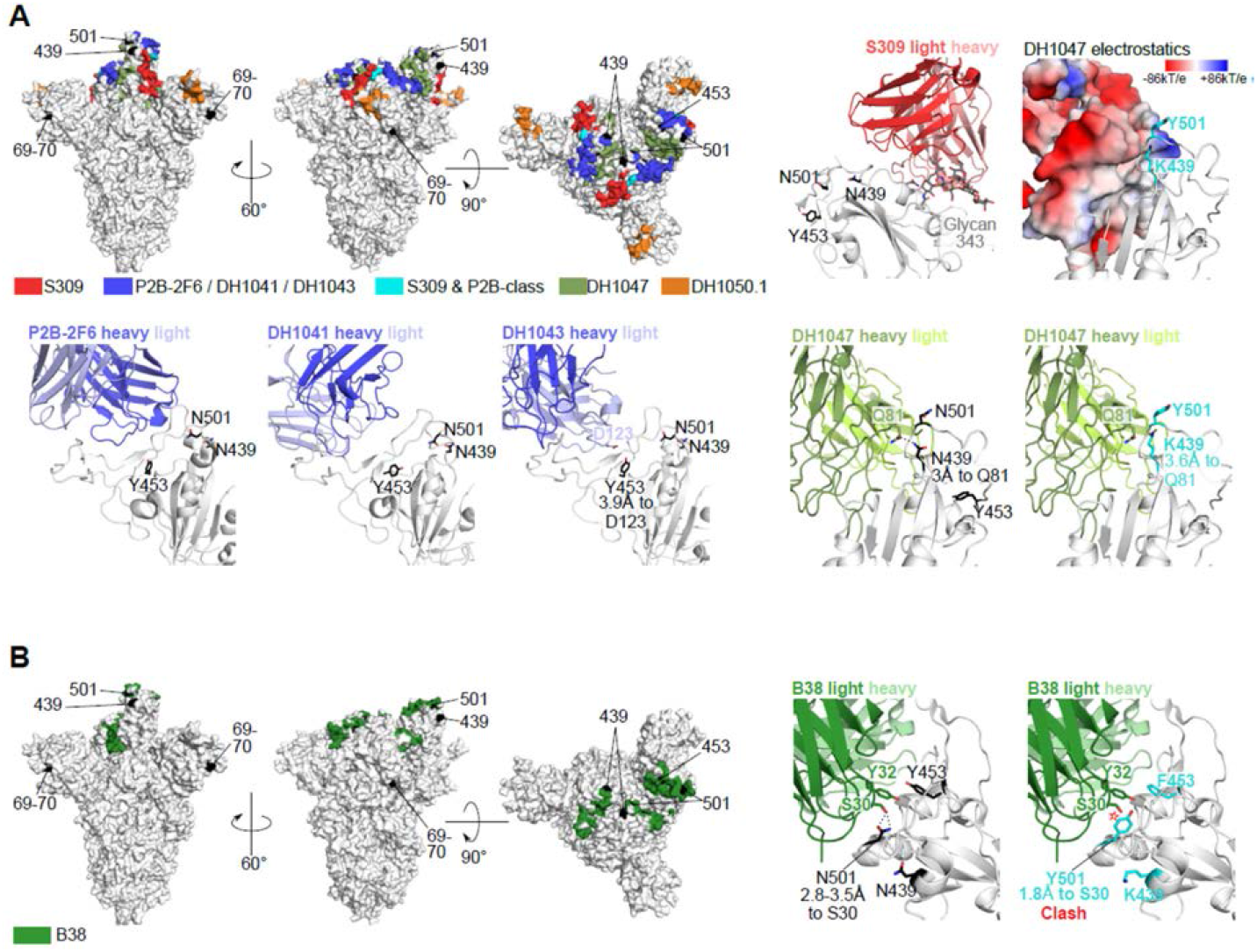
Structural analyses for antibody resistance mutations. (A) Top left 3 panels: Full spike trimer with antibody epitopes for S309, P2B-2F6, DH1041, DH1043, DH1047 and DH1050.1. Epitopes for P2B-2F6, DH1041 and DH1043 are similar and are grouped together. Top row, second from right panel shows location of spike sites 439, 453 and 501 with respect to S309. These spike sites are not close to S309 (>11Å). Top row rightmost panel shows the DH1047 antibody colored according to vacuum electrostatic potential, and the modeled mutations at spike sites Lys-439 and Tyr-501. Bottom row, rightmost two panels DH1047 interaction with sites 439, 453 and 501 using wildtype amino acids (second from right) and modeled mutations (rightmost). Bottom row, left 3 panels show the location of spike sites 439, 453 and 501 with P2B-2F6, DH1041 and DH1043. Polar interactions between antibody and spike residues of interest are shown with dotted black lines. (B) Similar to (A) except with B38 antibody. The modeled Tyr-501 is predicted to clash with light chain Ser-30 (~1.8Å, red star).

The decreased neutralization observed for S309 by ΔH69-V70 and N501Y could not be explained by structural analyses. Both the RBD mutations and ΔH69-V70 are distal from S309 (>11Å), suggesting allosteric interactions. Notably, it is the only mAb in this study that interacts strongly with a glycan (at site 343), and that changes in spike dynamics or conformations can impact glycan processing (34).

## DISCUSSION

Recent months have seen the emergence of a growing number of novel SARS-CoV-2 variants that can rapidly and repeatedly shift to prevalence in local populations and even globally (5, 35). Newer variants carry multiple Spike mutations (12–14) that are a potential concern for immune escape. The B.1.1.7 variant studied here was first detected in England in September 2020, where it rapidly came to dominate the regional pandemic and has now been detected in over 40 countries. Variants in the UK and Denmark followed a shifting dynamic, starting with the emergence of the G clade as the dominant form, followed by increasing prevalence of the GV clade that mirrored across Europe (35), and then regional appearance of variants that carried combinations of ΔH69-V70 with Y453F and N439K, finally to be followed by the introduction of the B.1.1.7 variant, which rapidly rose to dominance in the UK, and is now beginning to increase in frequency in Denmark. The serial waves of variant prevalence in these two countries suggest complex dynamics that may come into play as the SARS-CoV-2 continues to evolve. Furthermore, co-circulation of major variants in a geographically local region may enable recombination (36) bringing together mutations that enhance fitness either through infectivity or immunological resistance.

Prior to the emergence of this variant, two SARS-CoV-2 vaccines based on ancestral Spike proved highly effective and recently received emergency use authorization, including the Moderna mRNA-1273 vaccine studied here (9) and a similar mRNA vaccine developed by Pfizer/BioNTek (10). Another vaccine based on ancestral Spike nanoparticles developed by Novavax (37) is currently undergoing phase 3 testing in the UK, USA and Mexico, with phase 1/2 testing ongoing in South Africa. In addition to vaccines, several potent RBD-specific mAbs have received emergency use authorization for treatment of mild-to-moderate COVID-19 in the USA (FDA Press Release, November 21, 2020 and November 09, 2020), while still other therapeutic mAbs are in development (38) (also see recent announcement)

Here we show in a lentivirus-based pseudovirus assay that variant B.1.1.7 exhibits only modestly reduced neutralization susceptibility in the presence of convalescent sera (1.5-fold average) and sera from the Moderna and Novavax phase 1 studies (2-fold average after two inoculations) using the prototypic D614G variant as comparator. While it is not known for certain what level of neutralization is required for the remarkable efficacy in phase 3 studies completed to date, it is noteworthy that both the Moderna and Pfizer/BioNTech mRNA vaccines demonstrated substantial efficacy prior to the second (final) dose (9, 10). Neutralization titers have been shown to increase by approximately 10-fold after the second dose for both vaccines (39–41), suggesting that a 2-fold reduction in neutralization will have minimal impact on vaccine efficacy in people who receive both doses of vaccine. It may be prudent to receive the second dose in a timely manner in regions where the B.1.1.7 variant circulates.

In contrast to our findings with polyclonal sera from convalescent individuals and vaccine recipients, the B.1.1.7 variant exhibited markedly reduced susceptibility to a subset of RBD-specific mAbs. Partial escape from four mAbs (COVA1-18, COVA2-15, S309 and to a lesser extent, B38) was associated with the N501Y mutation. Modest escape from two additional mAbs (DH1042 and H4) could not be mapped with the specific mutations tested. Notably, B.1.1.7 exhibited no escape from four RBD-specific mAbs tested here (DH1041, DH1043, DH1047 and P2B-2F6).

In summary, our findings indicate that B.1.1.7 is not a neutralization escape variant of concern for vaccine efficacy and the risk of re-infection. In addition, although the variant is considerably less susceptible to certain mAbs, other RBD-specific mAb retain full activity. While this is encouraging, it is becoming increasingly clear that SARS-CoV-2 continues to evolve and that new variants may arise that pose a greater risk for immune escape. Early identification and characterization of newly emerging variants requires robust genetic surveillance coupled with rapid laboratory and clinical investigation to facilitate the timely design and testing of next generation vaccines and therapeutic mAbs should they be needed.

## ACKNOWLEDGEMENTS

We thank the HIV Vaccine Trials Network and HIV Prevention Trial Network for serum samples from COVID-19 convalescent individuals. We thank Peter Kwong for generously sharing the mAbs B38, H4, P2B-2F6, and S309 produced at the Vaccine Research Center, NIH. We also thank Jin Tong, Elize Domin, Wenhong Feng and Miroslawa Bilska for excellent technical assistance. Original data and specimens for Protocol 20-0003 were supported by the Division of Microbiology and Infectious Diseases, National Institute of Allergy and Infectious Diseases. KW and BK were supported by LANL LDRD 20190441ER. DCM, XS, HT, and CM were supported by the COVID-19 Network (CoVPN) and the National Institute of Health. DL, BFH, and KOS were supported by a grant from the State of North Carolina from federal CARES Act funds, and NIAID grant AI142596.

## AUTHOR CONTRIBUTIONS

DCM designed the study, coordinated assays, data analysis, and manuscript preparation, and helped write and edited the manuscript. XS helped with study design, coordinated the study, performed data analysis and visualization, and wrote the manuscript. HT participated in study design, site-directed mutagenesis, data generation, and manuscript writing and editing. CM participated in data generation and reviewing, and manuscript review and editing. DL and KOS produced, purified and provided mAbs and edited the manuscript. BFH provided mAbs and edited the manuscript. NK contributed data, data visualizations, and to manuscript preparation. BK helped with study design, generated data visualizations, and contributed to data interpretation and manuscript preparation. JT, HY, and WF also helped generate data visualizations. KW provided structural analyses and helped data interpretation and manuscript preparation..

## DECLARATION OF INTERESTS

Rolando Pajon is an employee of Moderna, Inc. Filip, Dubovsky, Gale Smith and Gregory M. Glenn are employees of Novavax, Inc.

## MATERIALS AND METHODS

### Ethics Statement

Clinical trials described in this manuscript were approved by the appropriate Institutional Review Boards (IRBs).

### Spike mutant production

Construction of full-length SARS-CoV-2 Spike with Wuhan-1 and the D614G variant sequences have been reported previously. Plasmids for B.1.1.7 and subvariants were generated by site-directly mutagenesis. Positive clones were fully sequences to ensure that all intended mutations were present and no additional mutations were introduced for each variant.

### Serum samples

Sera for the mRNA-1273 phase 1 study (NCT04283461) were obtained from the Division of Microbiology and Infectious Diseases, National Institute of Allergy and Infectious Diseases for the mRNA-1273 phase 1 study team and Moderna Inc. The phase 1 study protocols and results are reported previously (39, 41). The phase 1 trial tested the identical vaccine (mRNA-1273), dose (100μg) and schedule as used in the Moderna phase 3 (NCT04470427). mRNA-1273 is a lipid nanoparticle (LNP)-encapsulated mRNA-based vaccine that encodes for a full-length, prefusion stabilized spike protein of SARS-CoV-2 (9). Samples tested against the B.1.1.7 variant (together with D614G as control) were collected at day 29 (4 weeks post 1^st^ inoculation) or day 57 (4 weeks post 2^nd^ inoculation). Samples tested against the subvariants (together with D614G as control) were all from day 57.

Novavax phase 1 sera were obtained from Novavax. The phase 1 study (NCT04368988) tested a 5 μg dose of SARS-CoV-2 recombinant nanoparticle vaccine with or without Matrix-M adjuvant (37). Serum samples (N=28) tested here were from the vaccine arm with the Maxtrix-M adjuvant, which is the identical vaccine in the ongoing Novavax global phase 3 study (NC04611802). Samples tested were randomly collected and not pre-selected for higher titer respones at 2 weeks post 2^nd^ inoculation (day 35).

Convalescent sera were collected in an observational cohort study conducted by the HIV Vaccine Trial Network and the HIV Prevention Trials Network (protocol HVTN 405/HPTN 1901; NCT04403880). Samples were collected from the first visit of the study, scheduled at 1-8 weeks post resolution of COVID-19, or 2-10 weeks post most recent positive SARS-CoV-2 test, if asymptomatic. The subset of samples included in this study were pre-selected as representing high, medium and low neutralization titers against the D614G variant of SARS-CoV-2.

### MAbs

Antibodies B38, H4, P2B-2F6, and S309 (42–44), were provided by Dr. Peter Kwong. Antibodies DH1041, DH1042, DH1043, and DH1047 were provided by Drs. Kevin Saunders, Dapeng Li, and Barton Haynes (45). Antibodies COVA1-18 and COVA2-15 were provided by Dr. Rogier Sanders (46).

### Cells

HEK 293T/17 cells (ATCC cat. no. CRL-11268) and 293T/ACE2.MF (provided by Drs. Michael Farzan and Huihui Mu) were maintained in 12 mL of growth medium (DMEM, 10% heat-inactivated fetal bovine serum, 50 μg gentamicin/ml, 25mM HEPES) in T-75 culture flasks in a humidified 37°C, 5% CO2 environment. Puromycin (3 μg/mL) was added to the growth medium for maintaining 293T/ACE2.MF cells. Cells were split at confluency using TrypLE Select Enzyme solution (Thermo Fisher Scientific).

### Pseudotyped virus production

SARS-CoV-2 Spike-pseudotyped viruses were prepared and titrated for infectivity essentially as described previously (5). An expression plasmid encoding codon-optimized full-length Spike of the Wuhan-1 strain (VRC7480), was provided by Drs. Barney Graham and Kizzmekia Corbett at the Vaccine Research Center, National Institutes of Health (USA). Mutations were introduced into VRC7480 by site-directed mutagenesis using the QuikChange Lightning Site-Directed Mutagenesis Kit from Agilent Technologies (Catalog # 210518) using primers as listed in **Table S3**. All mutations were confirmed by full-length Spike gene sequencing. Pseudovirions were produced in HEK 293T/17 cells (ATCC cat. no. CRL-11268) by transfection using Fugene 6 (Promega Cat#E2692) and a combination of Spike plasmid, lentiviral backbone plasmid (pCMV ΔR8.2) and firefly Luc reporter gene plasmid (pHR′ CMV Luc) (47) in a 1:17:17 ratio in Opti-MEM (Life Technologies). Transfection mixtures were added to pre-seeded HEK 293T/17 cells in T-75 flasks containing 12 ml of growth medium and incubated for 16-20 hours at 37°C. Medium was removed and 15 ml of fresh growth medium added. Pseudovirus-containing culture medium was collected after an additional 2 days of incubation and clarified of cells by low-speed centrifugation and 0.45 μm micron filtration.

TCID50 assays were performed prior to freezing aliquots of the viruses at −80°C. Viruses were serially diluted 3-fold or 5-fold in quadruplicate for a total of 11 dilutions in 96-well flat-bottom poly-L-lysine-coated culture plates (Corning Biocoat). An additional 4 wells served as background controls; these wells received cells but no virus. Freshly suspended 293T/ACE2.MF cells were added (10,000 cells/well) and incubated for 66-72 hours. Medium was removed by gentle aspiration and 30 μl of Promega 1X lysis buffer was added to all wells. After a 10 minute incubation at room temperature, 100 μl of Bright-Glo luciferase reagent was added to all wells, mixed, and 105 μl of the mixture was added to a black/white plate (Perkin-Elmer). Luminescence was measured using a GloMax Navigator luminometer (Promega). TCID50 was calculated using the method of Reed and Muench as described (48). HIV-1 p24 content (produced by the backbone vector) was quantified using the Alliance p24 ELISA Kit (PerkinElmer Health Sciences, Cat# NEK050B001KT). Relative luminescence units (RLUs) were adjusted for p24 content.

### Neutralization assay

Neutralization was measured in a formally validated assay that utilized lentiviral particles pseudotyped with SARS-CoV-2 Spike and containing a firefly luciferase (Luc) reporter gene for quantitative measurements of infection by relative luminescence units (RLU). A pre-titrated dose of virus was incubated with 8 serial 5-fold dilutions of serum samples in duplicate in a total volume of 150 μl for 1 hr at 37°C in 96-well flat-bottom poly-L-lysine-coated culture plates. Cells were detached using TrypLE Select Enzyme solution, suspended in growth medium (100,000 cells/ml) and immediately added to all wells (10,000 cells in 100 μL of growth medium per well). One set of 8 wells received cells + virus (virus control) and another set of 8 wells received cells only (background control). After 66-72 hrs of incubation, medium was removed by gentle aspiration and 30 μl of Promega 1X lysis buffer was added to all wells. After a 10 minute incubation at room temperature, 100 μl of Bright-Glo luciferase reagent was added to all wells. After 1-2 minutes, 110 μl of the cell lysate was transferred to a black/white plate. Luminescence was measured using a GloMax Navigator luminometer (Promega). Neutralization titers are the inhibitory dilution (ID) of serum samples, or the inhibitory concentration (IC) of mAbs at which RLUs were reduced by either 50% (ID50/IC50) or 80% (ID80/IC80) compared to virus control wells after subtraction of background RLUs. Serum samples were heat-inactivated for 30 minutes at 56°C prior to assay.

### Phylogenetic trees

The tree in Fig. S1 is based on the GISAID data sampled on Jan. 17th, 2021, and passed through a quality control filter, and presented as the “tree of the day” at the cov.lanl.gov website https://cov.lanl.gov/components/sequence/COV/rainbow.comp. The “full” alignment was used as described previously (5). Only the mutations of interest for this study are tracked in this tree for clarity. The tree is rooted using the Wuhan-Hu-1 isolate (GenBank accession NC_045512).

All Phylogenetic trees are constructed using parsimony, TNT version 1.5 (49), with 5 or 10 random-sequence addition replicates with TBR (tree-bisection-reconnection) branch swapping (command: “mult = rep REPS tbr hold 1 wclu 1000”, where REPS equals 5 or 10, with the bbreak cluster value set to 40).

### Structural analyses

We used PDB: 7C2L (50) for the full trimeric spike structure, and antibody spike complex structures from Li et al. (45) for DH1041-DH1047 antibodies, PDB: 6WPS (44) for S309, PDB: 7BWJ (43) for P2B-2F6, and PDB: 6XDG for REGN antibodies (51). Antibody epitopes were defined as spike amino acids with any heavy atoms within 4Å antibody heavy atoms (33). Antibody epitope, electrostatics and polar bonds calculations as well as mutation modeling were performed in PyMOL (The PyMOL Molecular Graphics System, Version 2.0 Schrödinger, LLC.). Mutations were modeled with spike or RBD in isolation and the rotamers with least predicted strain in PyMOL were used. For B38, to identify the most amenable Y-501 rotamers, the N501Y mutation was modeled with the antibody-RBD complex; however, all identified rotamers induced substantial clashes and the rotamer with the least clash was retained. PyMOL was also used for structural renderings for all figures.

### Statistical analyses

Neutralization ID50 titers or IC50 concentrations between each variant and D614G, or between other pairs of variants with and without the N501Y or ΔH69-V70 were compared using the Wilcoxon signed-rank 2-tailed test. Decrease in ID50 and ID80 titer for Moderna Day 29 and Day 57 samples against B.1.1.7 were compared using the Wilcoxon rank-sum test, 2-tailed. To correct for multiple test corrections, false discovery rates (FDR or q values) were calculated as in (52) implemented in a Python package (https://github.com/nfusi/qvalue) for Python version 3.4.2. All tests with q < 0.1 were considered as significant, which corresponded to p < 0.042. Wilcoxon signed-rank test and Wilcoxon rank-sum test were performed using the coin package (version 1.3-1) with R (version 3.6.1). Wilcoxon signed-rank test and Wilcoxon rank-sum test were performed using the coin package (version 1.3-1) with R (version 3.6.1).

## Supplemental Materials

**Fig. S1.**
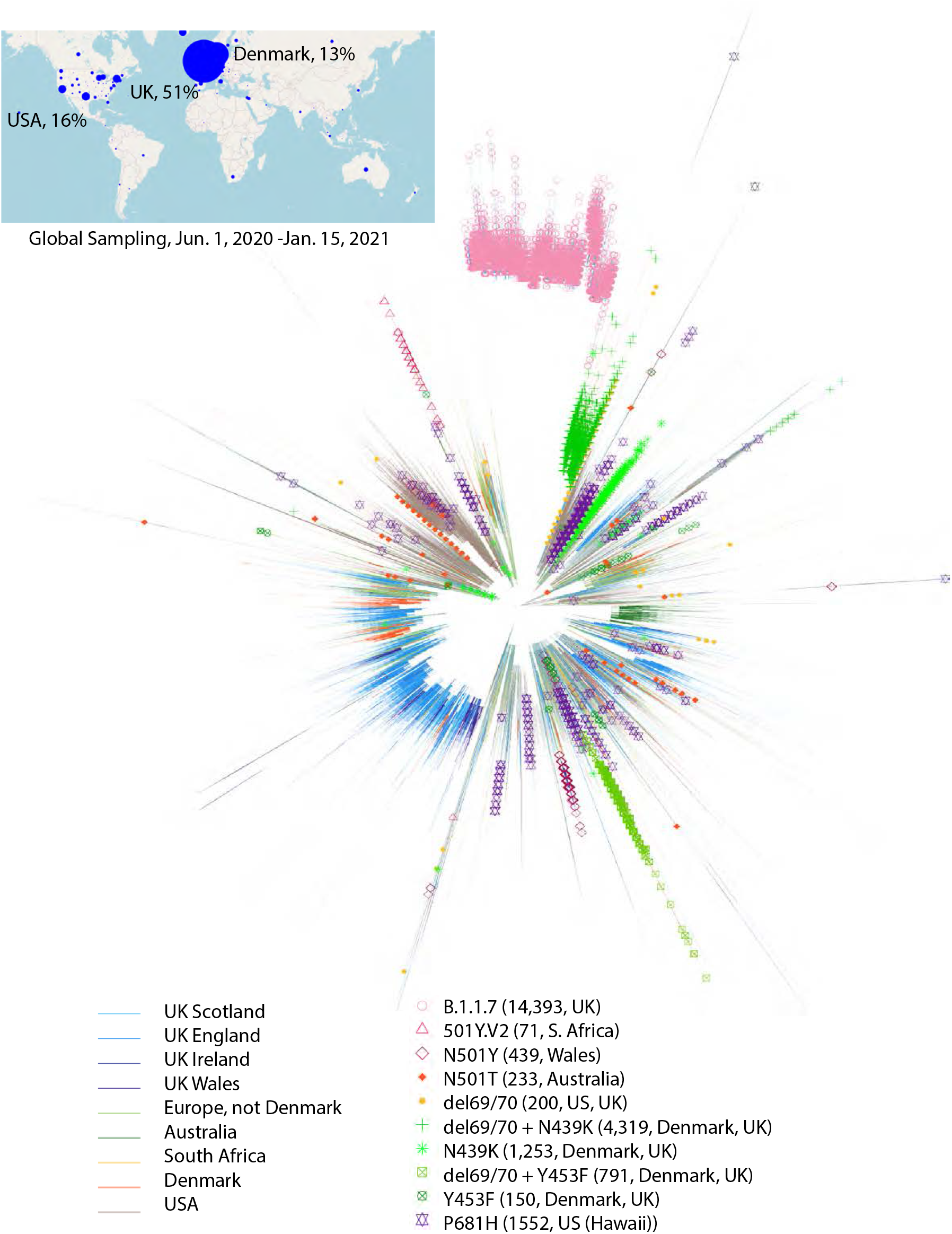
Parsimony tree showing the relationships between B.1.1.7 and other variants with key mutations. 244,291 seqs were included in this tree, which is based on the GISAID data sampled on Jan.17th, 2021. Countries where the variants of interest are commonly found are indicated by branch color, key variants by symbols at the tips. The B.1.1.7 lineage is shown as pink open circles. The B.1.1.7 variant is still predominantly found in the UK, but by Jan. 24^th^ 2021 had been sampled in 44 countries, and in 20 different states in the USA. Some of these geographically diverse samples were likely to have been detected as a consequence of sampling travelers from the UK and their contacts, due to the international interest in B.1.1.7 in late Dec. 2020 and early Jan. 2021; regardless of this potential sampling bias, the B.1.1.7 variant clearly has a global presence. The N501Y mutations is also found in the South African Variant of Interest, 501Y.V2, the red open triangles. N501Y has also transiently but significantly emerged in local populations, as a lone Spike mutation the D614G background, found in the early summer in Victoria, Australia, and also found in Wales (in this figure, and followed over time in Fig. 1). A distinct variant, N501T as been emerging in Sydney, New South Wales, Australia, that has become increasing common through December, 2020. Variants carrying the only the deletion in Spike at Δ69/70 on a D614G background; on the rare occasions they are found, they were primarily sampled in the UK and US. In contrast, Δ69/70 is frequently found coupled with other RBM mutations, either embedded in B.1.1.7 or coupled with N439K or Y453F, commonly circulating in both Denmark and UK as well as in other European counties; N439K and Y453F variants not coupled to Δ69/70 are found less frequently (see Fig. 1B). Of note, P681H is embedded in B.1.1.7, but is also frequently found independently, and is often found to increase in frequency regionally when it does arise, and it is recurrent throughout the phylogeny; it is the dominant form in the Hawaiian epidemic, and is found in many places throughout the US. All of the currently variants of interest are in the G clade, Spike D614G background. The map insert includes the subset of complete GISAID sequences with a known sampling date that were sampled since June 1, 2020, and the area of the circle is proportional to sample size, illustrating the strong sampling bias in the data: the UK is contributing over half of the sequences in the dataset, with the USA and Denmark following, these three nations account for 80% of the GISAID sequences that pass through the cov.lanl.gov quality control screen.

**Fig. 2.**
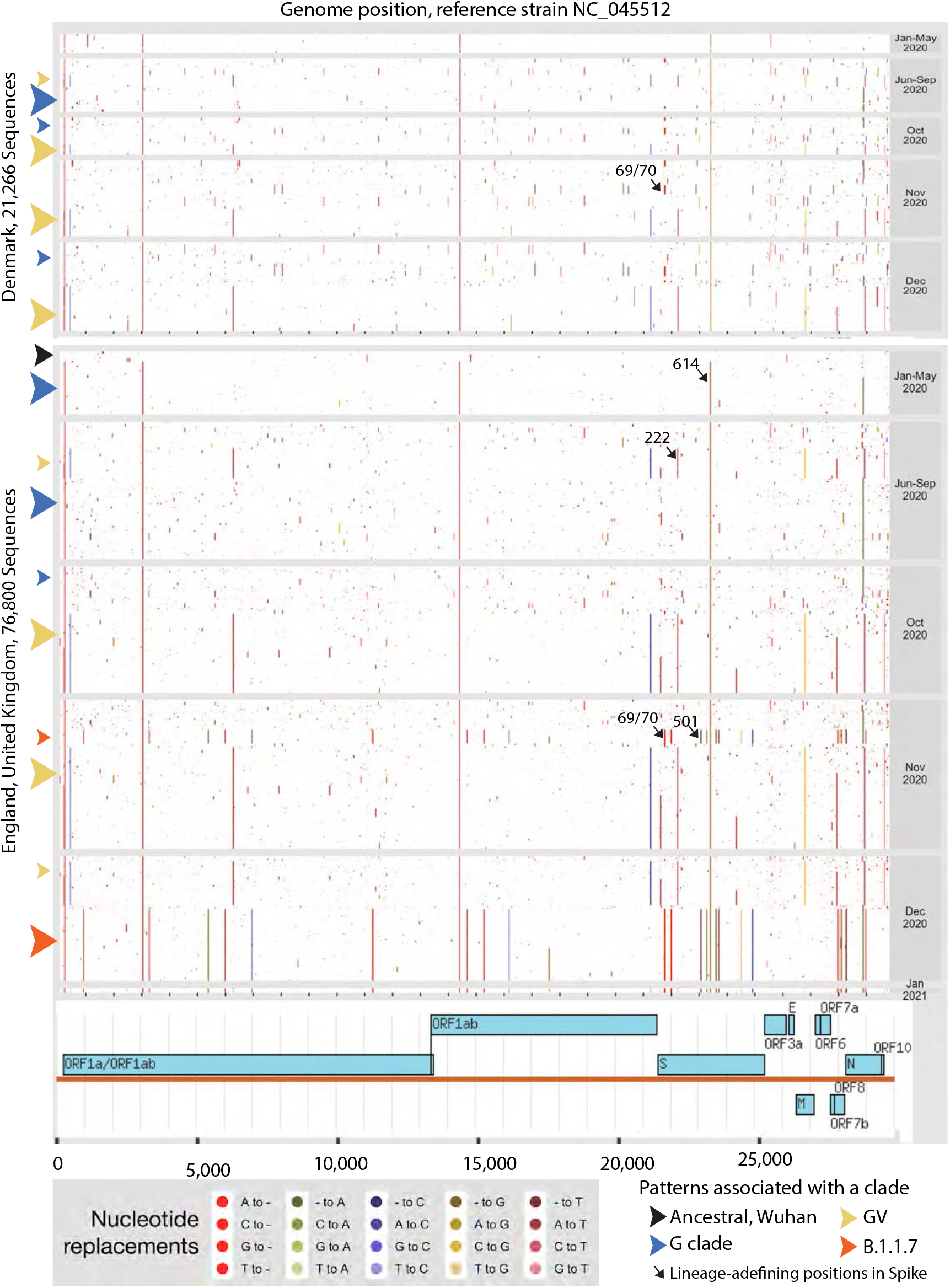
Highlighter plot showing the mutations associated with major clade in the UK and Denmark transitioning over time. Distribution of mutations among SARS-CoV-2 sequences from Denmark and from England, UK. These figures are highly compressed pixel plots representing the full sampling of complete sequences from these countries, nearly 100,000 sequence in all, and full length genomes across the x-axis. Each row in the matrix represents a single genome sequence; each column, a genome position. Colored dots denote locations of nucleotide mutations relative to the Wuhan-Hu-1 isolate (GenBank accession NC_045512); unmutated bases are white to allow visualization of mutations. Sequences are presented in separate panels based on time of sampling. Within each panel, sequences are clustered phylogenetically, ordered top-to-bottom by position within a parsimony-based phylogenetic tree; therefore, mutations at a particular locus that are shared within a lineage are seen as vertical lines, and groupings of vertical lines that start and end at the same heights indicate coherent lineages defined by multiple mutations. Clades with more members are sorted lower. Colored arrowheads at the left margin indicate lineages of particular interest; smaller and larger arrowheads reflect the proportional representation of lineages among samples from a particular time window.

**Fig. S3.**
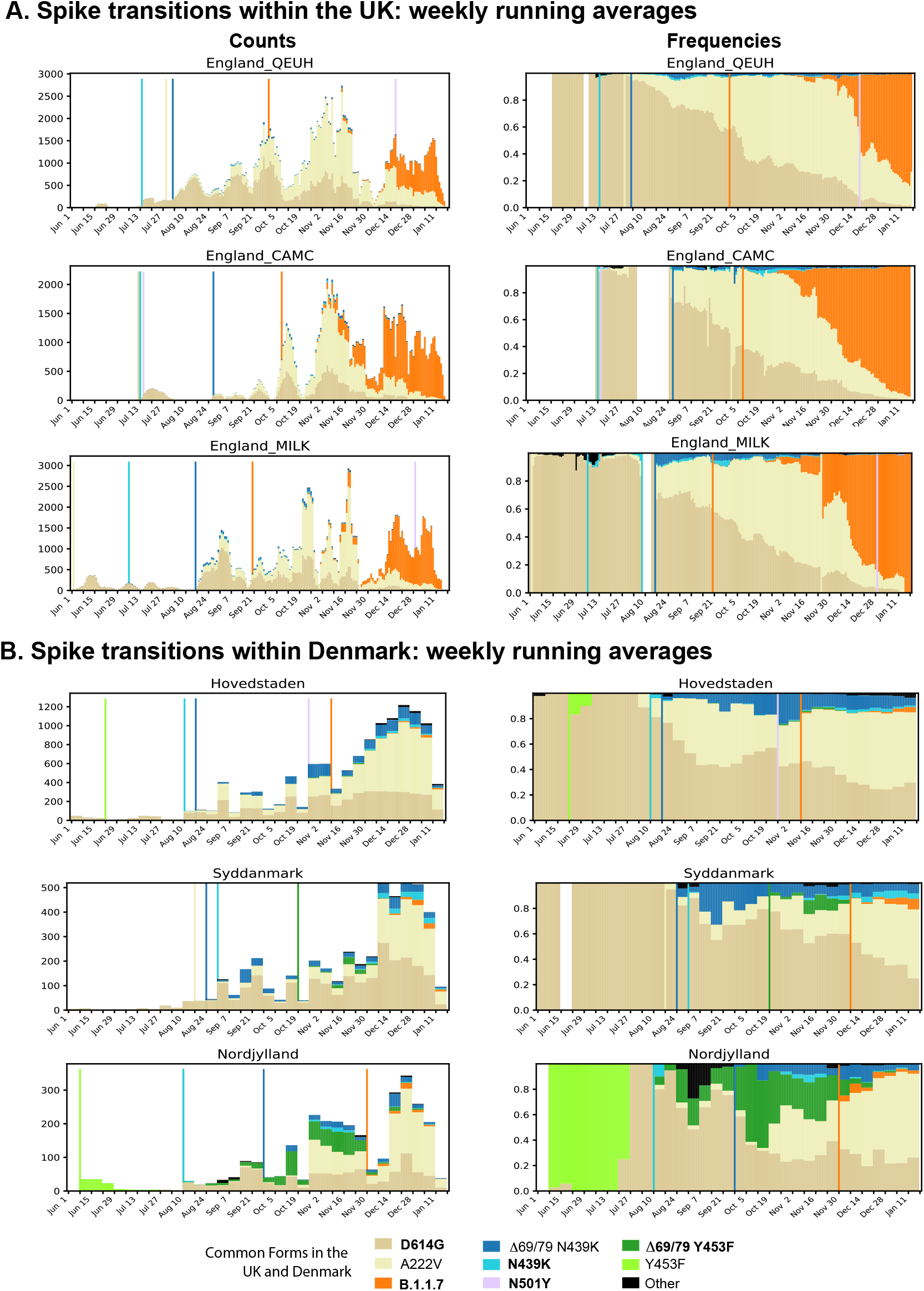
Transitions in key mutational patterns over time in local regions in Denmark and in England. These figures are drawn as in Fig. 1C. Weekly running averages are plotted each day, with the actual counts on the left, and relative frequencies on the right. Regions with white banks indicate no sampling was done in that time frame. **A. English regional data over time**. The overall pattern seen in England in Fig. 1 was consistent across more local sampling in England (QUEH, CAMC, and MILK). The increased frequency of the GV clade, carrying the 4 G clade mutations plus an additional 8 mutations including the A222V mutation begins in early August, and it increases in prevalence relative to the G clade in each local region, but the transitions are more gradual than the later transition to the B.1.1.7 form indicated in orange. The lines indicate when a particular variant was introduced into the region, and the orange line indicates the first introduction of B.1.1.7, first sampled in late September and early October in each region. After the B.1.1.7 variant is first introduced, a very gradual increase in frequency is observed for a period, and then it begins to appreciably increase in frequency. **B. Danish regional data over time**. The B.1.1.7 form was introduced into Danish cites in November/December, and by mid-Jan was becoming more consistently observed. Danish sampling times are indicated on a weekly basis, UK daily, hence the broader bands. The N439K and N453Y variants, particularly in combination with the Δ69/70 deletion, were more prominent among Danish samples than among the UK samples; both these and the original D614G form becoming less frequently sampled, and the GV form more prominent, at the point when the B.1.1.7 form was introduced into Denmark.

GV defining mutations from figure **S2**. 12 mutations, include the following G clade mutations are highlighted

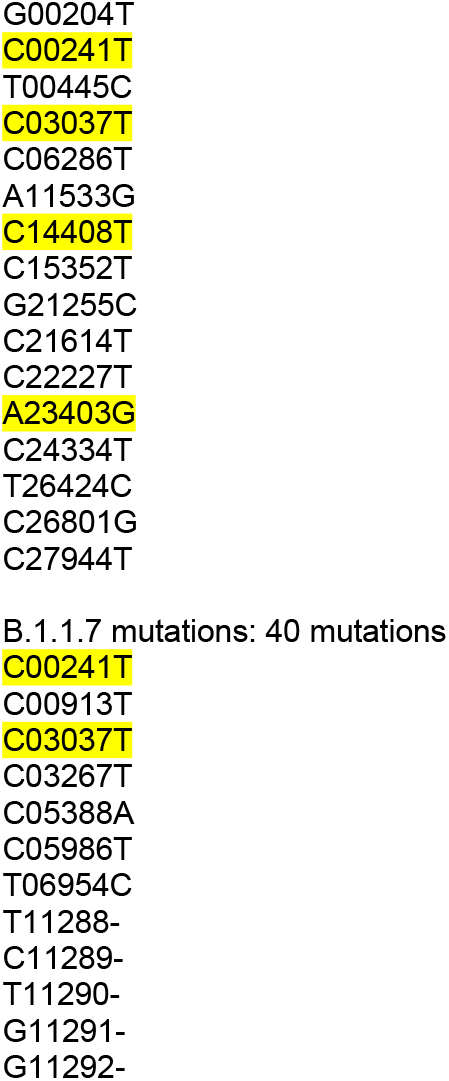
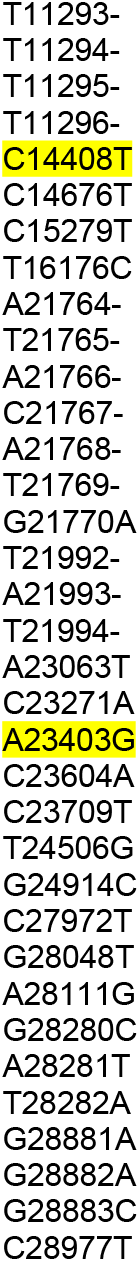

---

**Table S1.**
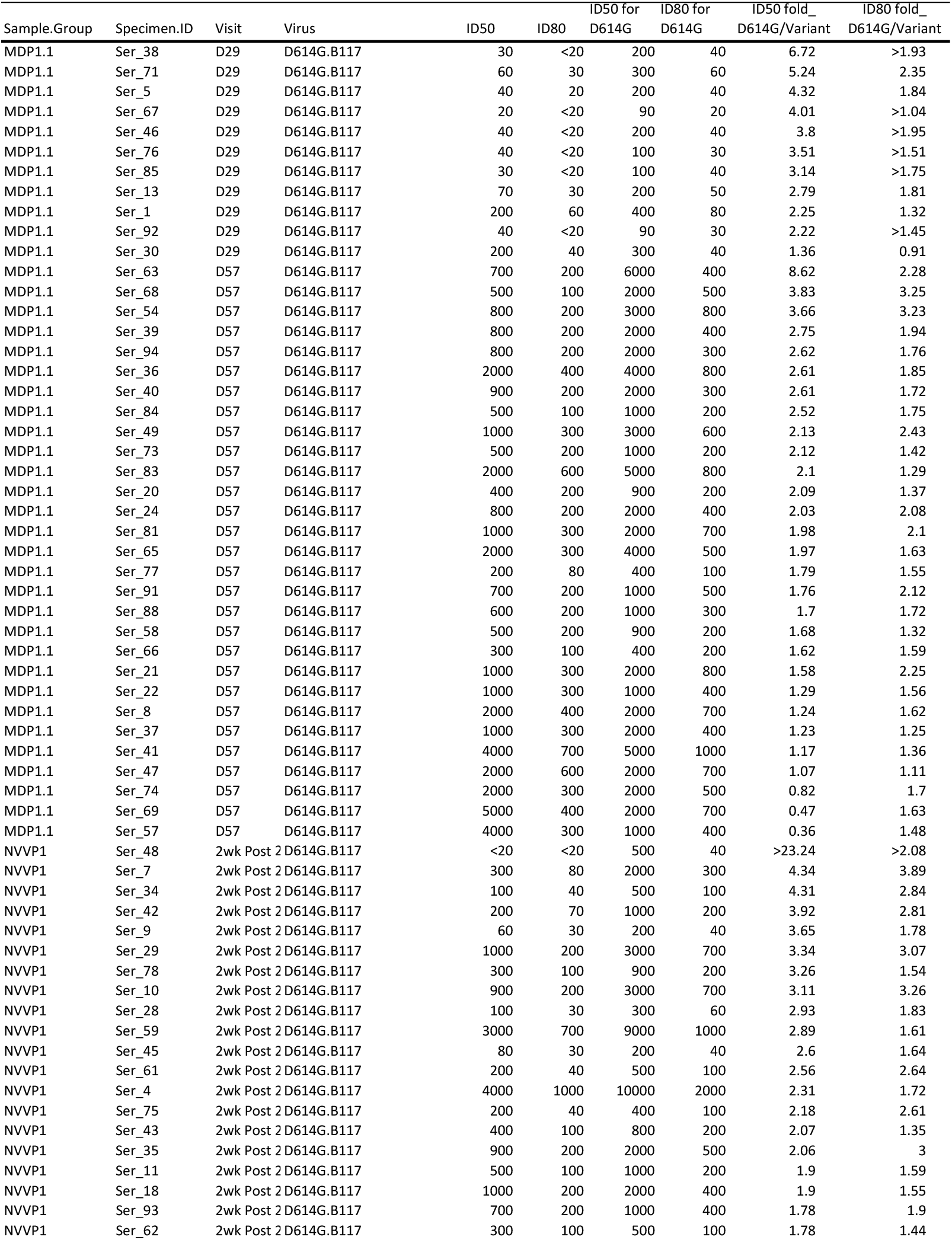
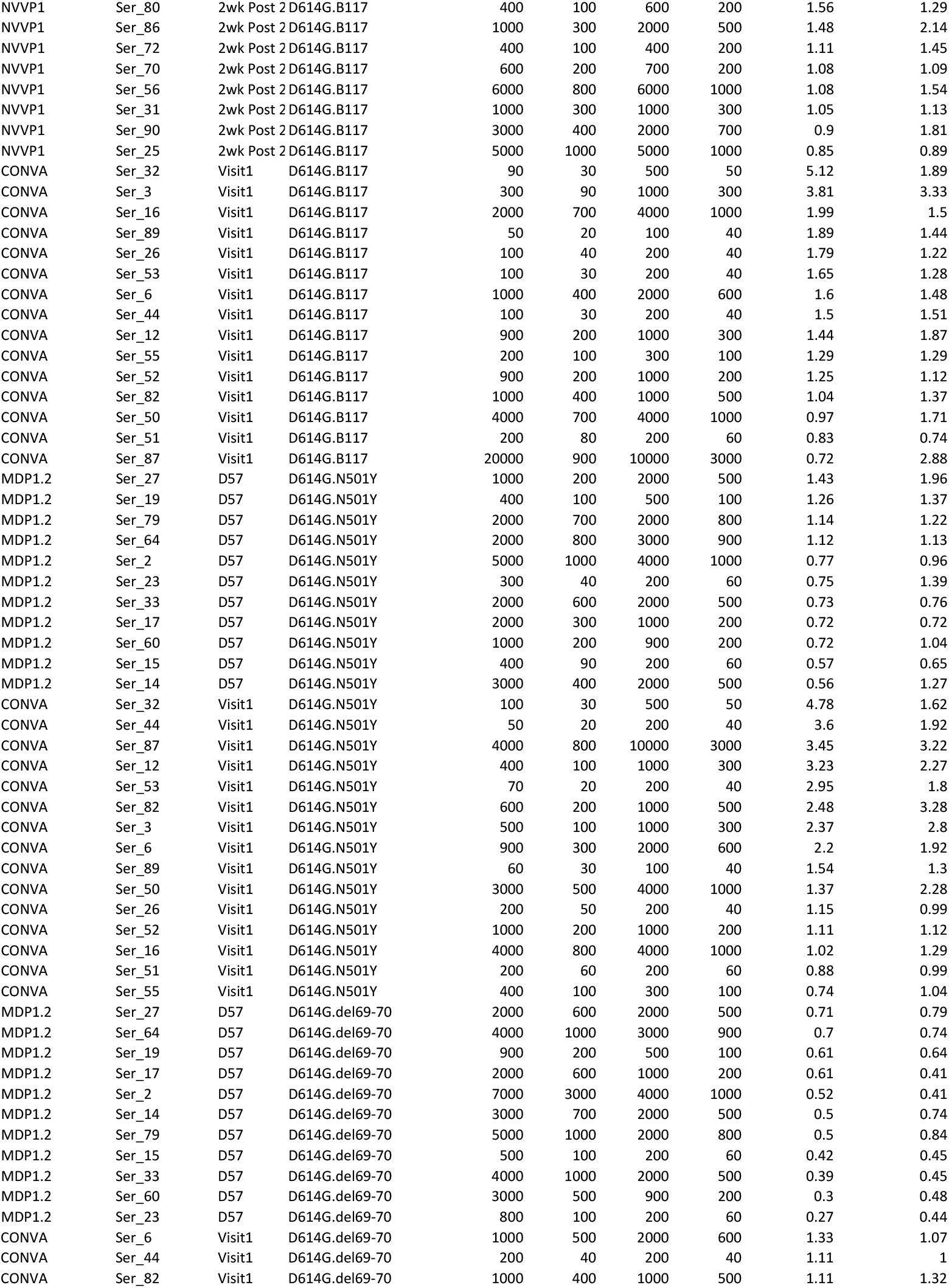
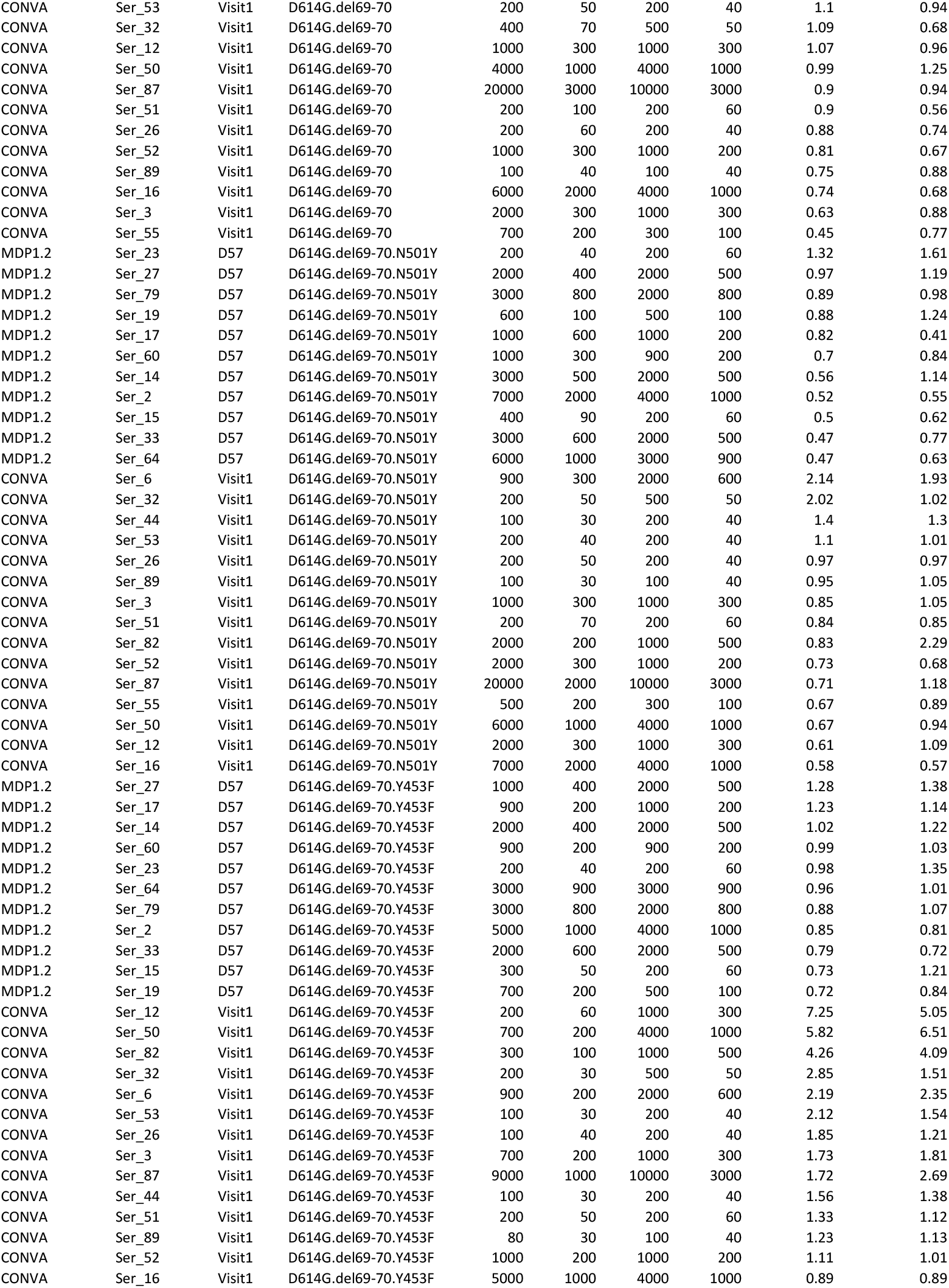
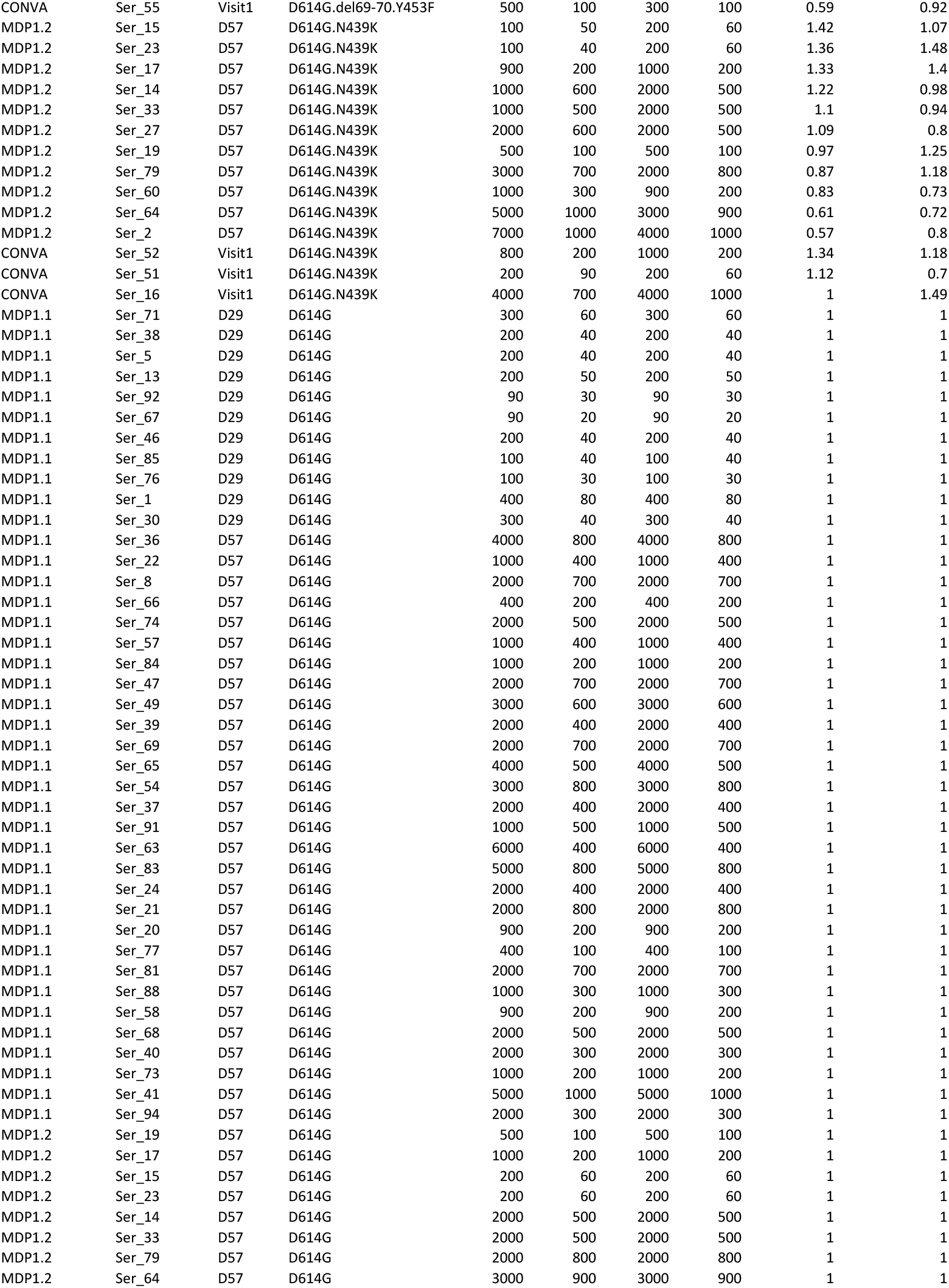
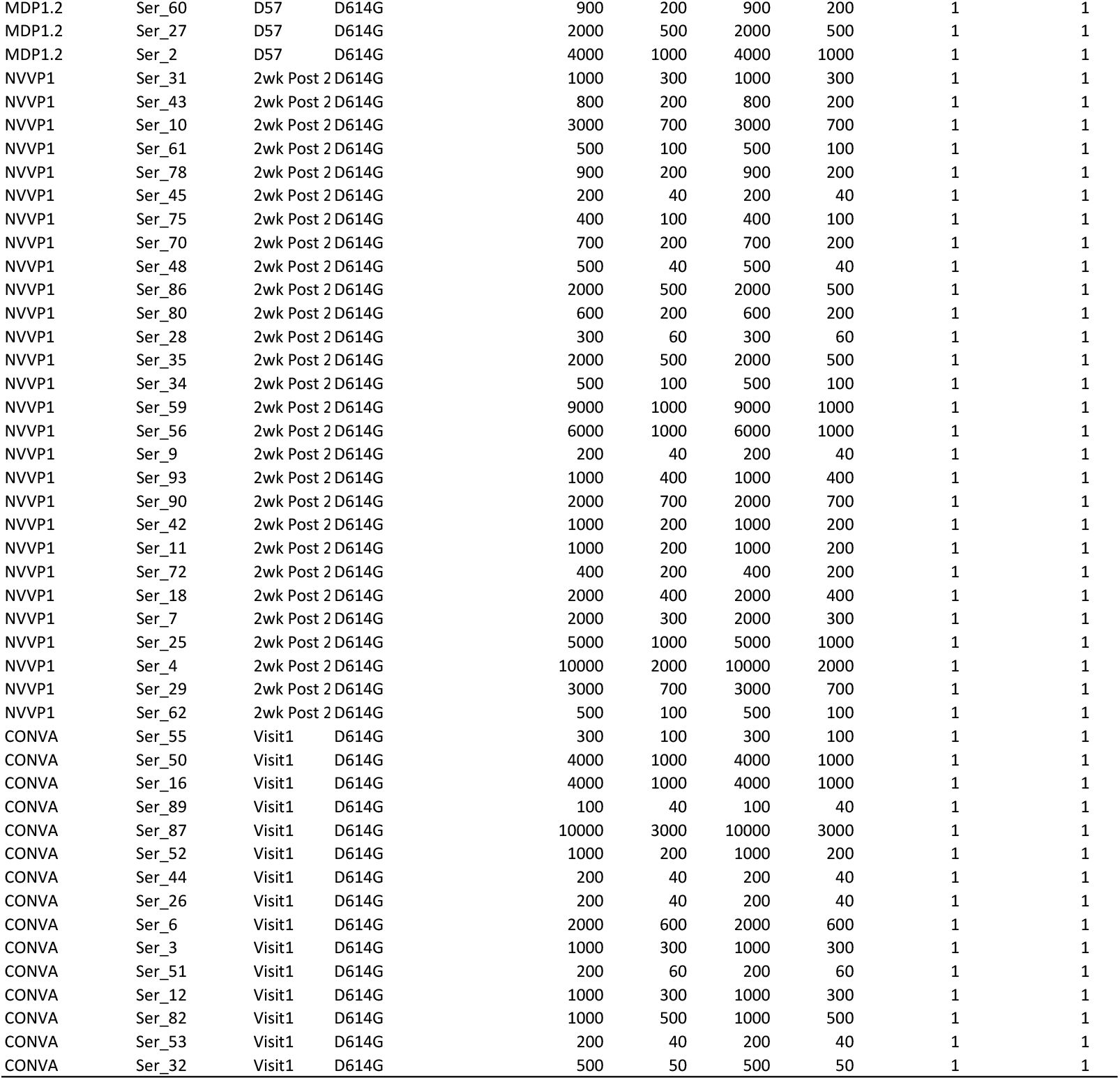
Serum neutralization titers and fold difference (D6I4G/Variant).

**Table S2.** Statistical results.

| Variable 1 | Variable 2 | Group Identifier | Measurement | Sample Size | p value | q value | median_<br>Variable 1 | median_<br>variable 2 | Figure<br>referece | Test |
| --- | --- | --- | --- | --- | --- | --- | --- | --- | --- | --- |
| D614G | D614G.B117 | MDP1.1 | ID50 | 40 | **<u>5.85E-06</u>** | **<u>7.80E-05</u>** | 1374 | 687 | 2A | Wilcoxon signed-rank, paired, 2 tailed |
| D614G | D614G.B117 | NVVP1 | ID50 | 28 | **<u>1.77E-05</u>** | **<u>1.77E-04</u>** | 975 | 448 | 2A | Wilcoxon signed-rank, paired, 2 tailed |
| D614G | D614G.B117 | Convalescent | ID50 | 15 | 0.04126 | 0.069987 | 1105 | 325 | 2A | Wilcoxon signed-rank, paired, 2 tailed |
| D614G | D614G.B117 | MDP1.1 | ID80 | 40 | **<u>5.46E-12</u>** | **<u>2.18E-10</u>** | 347 | 188 | 2B | Wilcoxon signed-rank, paired, 2 tailed |
| D614G | D614G.B117 | NVVP1 | ID80 | 28 | **<u>1.54E-06</u>** | **<u>3.08E-05</u>** | 209 | 142 | 2B | Wilcoxon signed-rank, paired, 2 tailed |
| D614G | D614G.B117 | Convalescent | ID80 | 15 | <u>0.00061</u> | <u>0.002413</u> | 218 | 106 | 2B | Wilcoxon signed-rank, paired, 2 tailed |
| D614G | D614G.del69-70 | MDP1.2 | ID50 | 11 | <u>0.000977</u> | <u>0.002604</u> | 1538 | 2896 | 2A | Wilcoxon signed-rank, paired, 2 tailed |
| D614G | D614G.del69-70 | Convalescent | ID50 | 15 | 0.25238 | 0.315475 | 1105 | 1244 | 2A | Wilcoxon signed-rank, paired, 2 tailed |
| D614G | D614G.del69-70 | MDP1.2 | ID80 | 11 | <u>0.000977</u> | <u>0.002604</u> | 453 | 617 | 2B | Wilcoxon signed-rank, paired, 2 tailed |
| D614G | D614G.del69-70 | Convalescent | ID80 | 15 | 0.151428 | 0.224338 | 218 | 301 | 2B | Wilcoxon signed-rank, paired, 2 tailed |
| D614G | D614G.del69-70.N501Y | MDP1.2 | ID50 | 11 | 0.001953 | 0.004596 | 1538 | 1804 | 2A | Wilcoxon signed-rank, paired, 2 tailed |
| D614G | D614G.del69-70.N501Y | Convalescent | ID50 | 15 | 0.072998 | 0.112305 | 1105 | 923 | 2A | Wilcoxon signed-rank, paired, 2 tailed |
| D614G | D614G.del69-70.N501Y | MDP1.2 | ID80 | 11 | 0.206055 | 0.294364 | 453 | 475 | 2B | Wilcoxon signed-rank, paired, 2 tailed |
| D614G | D614G.del69-70.N501Y | Convalescent | ID80 | 15 | 0.719727 | 0.738181 | 218 | 219 | 2B | Wilcoxon signed-rank, paired, 2 tailed |
| D614G | D614G.del69-70.Y453F | MDP1.2 | ID50 | 11 | 0.240234 | 0.315475 | 1538 | 1365 | 2A | Wilcoxon signed-rank, paired, 2 tailed |
| D614G | D614G.del69-70.Y453F | Convalescent | ID50 | 15 | 0.012451 | 0.026042 | 1105 | 343 | 2A | Wilcoxon signed-rank, paired, 2 tailed |
| D614G | D614G.del69-70.Y453F | MDP1.2 | ID80 | 11 | 0.577148 | 0.616377 | 453 | 351 | 2B | Wilcoxon signed-rank, paired, 2 tailed |
| D614G | D614G.del69-70.Y453F | Convalescent | ID80 | 15 | 0.008362 | 0.018582 | 218 | 122 | 2B | Wilcoxon signed-rank, paired, 2 tailed |
| D614G | D614G.N439K | MDP1.2 | ID50 | 11 | 0.764648 | 0.764648 | 1538 | 1312 | 2A | Wilcoxon signed-rank, paired, 2 tailed |
| D614G | D614G.N439K | Convalescent | ID50 | 15 | 0.25 | 0.315475 | 1105 | 822 | 2A | Wilcoxon signed-rank, paired, 2 tailed |
| D614G | D614G.N439K | MDP1.2 | ID80 | 11 | 0.464844 | 0.546875 | 453 | 481 | 2B | Wilcoxon signed-rank, paired, 2 tailed |
| D614G | D614G.N439K | Convalescent | ID80 | 15 | 0.5 | 0.571429 | 218 | 185 | 2B | Wilcoxon signed-rank, paired, 2 tailed |
| D614G | D614G.N501Y | MDP1.2 | ID50 | 11 | 0.240234 | 0.315475 | 1538 | 1604 | 2A | Wilcoxon signed-rank, paired, 2 tailed |
| D614G | D614G.N501Y | Convalescent | ID50 | 15 | 0.00116 | 0.002899 | 1105 | 412 | 2A | Wilcoxon signed-rank, paired, 2 tailed |
| D614G | D614G.N501Y | MDP1.2 | ID80 | 11 | 0.464844 | 0.546875 | 453 | 347 | 2B | Wilcoxon signed-rank, paired, 2 tailed |
| D614G | D614G.N501Y | Convalescent | ID80 | 15 | <u>0.000305</u> | <u>0.001356</u> | 218 | 128 | 2B | Wilcoxon signed-rank, paired, 2 tailed |
| D614G.del69-70 | D614G.del69-70.N501Y | MDP1.2 | ID50 | 11 | 0.041992 | 0.069987 | 2896 | 1804 | 2A | Wilcoxon signed-rank, paired, 2 tailed |
| D614G.del69-70 | D614G.del69-70.N501Y | Convalescent | ID50 | 15 | 0.524475 | 0.58275 | 1244 | 923 | 2A | Wilcoxon signed-rank, paired, 2 tailed |
| D614G.del69-70 | D614G.del69-70.N501Y | MDP1.2 | ID80 | 11 | 0.013672 | 0.026042 | 617 | 475 | 2B | Wilcoxon signed-rank, paired, 2 tailed |
| D614G.del69-70 | D614G.del69-70.N501Y | Convalescent | ID80 | 15 | 0.063721 | 0.101953 | 301 | 219 | 2B | Wilcoxon signed-rank, paired, 2 tailed |
| D614G.del69-70 | D614G.del69-70.Y453F | MDP1.2 | ID50 | 11 | <u>0.000977</u> | <u>0.002604</u> | 2896 | 1365 | 2A | Wilcoxon signed-rank, paired, 2 tailed |
| D614G.del69-70 | D614G.del69-70.Y453F | Convalescent | ID50 | 15 | **<u>6.10E-05</u>** | **<u>3.49E-04</u>** | 1244 | 343 | 2A | Wilcoxon signed-rank, paired, 2 tailed |
| D614G.del69-70 | D614G.del69-70.Y453F | MDP1.2 | ID80 | 11 | <u>0.000977</u> | <u>0.002604</u> | 617 | 351 | 2B | Wilcoxon signed-rank, paired, 2 tailed |
| D614G.del69-70 | D614G.del69-70.Y453F | Convalescent | ID80 | 15 | **<u>6.10E-05</u>** | **<u>3.49E-04</u>** | 301 | 122 | 2B | Wilcoxon signed-rank, paired, 2 tailed |
| D614G.N501Y | D614G.del69-70.N501Y | MDP1.2 | ID50 | 11 | 0.041992 | 0.069987 | 1604 | 1804 | 2A | Wilcoxon signed-rank, paired, 2 tailed |
| D614G.N501Y | D614G.del69-70.N501Y | Convalescent | ID50 | 15 | **<u>6.10E-05</u>** | **<u>3.49E-04</u>** | 412 | 923 | 2A | Wilcoxon signed-rank, paired, 2 tailed |
| D614G.N501Y | D614G.del69-70.N501Y | MDP1.2 | ID80 | 11 | 0.013672 | 0.026042 | 347 | 475 | 2B | Wilcoxon signed-rank, paired, 2 tailed |
| D614G.N501Y | D614G.del69-70.N501Y | Convalescent | ID80 | 15 | <u>0.000183</u> | <u>0.000916</u> | 128 | 219 | 2B | Wilcoxon signed-rank, paired, 2 tailed |
| Moderna D29 | Moderna D57 | B117 | D614G/B117_ID | 11, 29 | <u>0.000664</u> | <u>0.002413</u> | 3.51 | 1.97 | 2C | Wilcoxon rank sum, 2 tailed |
| Moderna D29 | Moderna D57 | B117 | D614G/B117_ID | 11, 29 | 0.585559 | 0.616377 | 1.75 | 1.7 | 2C | Wilcoxon rank sum, 2 tailed |

Grey shade: fold difference between median of two variables in comparison <1.3 and >0.77 (reflecting <30% difference in median values).
Underlined: p<0.01
Bold and underlined: p<0.001

**Table S3.**
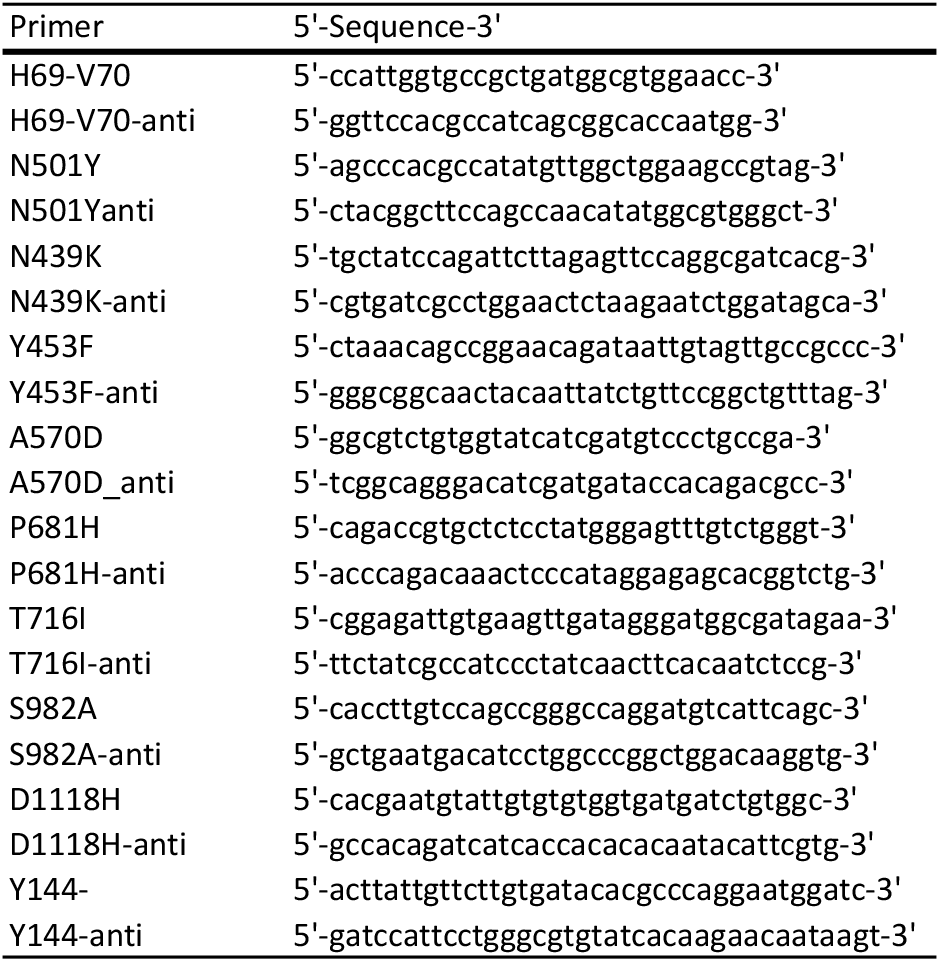
Primers used for site-directed-mutagenesis.

